# Mid-infrared free-electron laser-evoked discharge of crayfish compound eyes

**DOI:** 10.1101/131813

**Authors:** Fumio Shishikura, Heishun Zen, Ken Hayakawa, Yoshimasa Komatsuzaki, Yashushi Hayakawa, Takeshi Sakai, Toshiteru Kii, Hideaki Ohgaki

## Abstract

Light perception is an intriguing subject, and it has been demonstrated that some animals can perceive wavelengths beyond human vision. However, it is still controversial whether animals can see mid-infrared radiation. A combination of two free-electron laser (FEL) facilities, LEBRA-FEL (Nihon University) and KU-FEL (Kyoto University), can provide light sources with high peak power and spatially coherent monochromatic wavelengths that are continuously tuneable from 400 nm to 20 μm. We show that the crayfish (*Procambarus clarkii*) compound eye responds to pulsed mid-infrared FELs (1 Hz for KU-FEL; 2 Hz for LEBRA-FEL) from 3 to 17 μm. Our finding provides insight into the animal’s visual sensing of mid-infrared radiation, and that the sensing does not depend on thermal effects. Although the behavioural reason for vision in this wavelength range and the mechanism are still under investigation, understanding this type of visual sensing may lead to applications other than photophysical applications.

## INTRODUCTION

Compound eye vision is highly evolved in arthropods (Nilsson and Kelber, 2007; Yong, 2016) and is different from human vision. Arthropods see near-infrared radiation (Lindstrom and Meyer-Rochow, 1987; Schmitz et al., 2000) and ultraviolet light (El-Bakary and Sayed, 2011) in addition to visible light (Kennedy and Bruno, 1961). Crayfish, a highly evolved arthropod, have been investigated for 130 years since they were introduced to zoological laboratories (Huxley, 1880) and have contributed to the fundamental theory of vision physiology (Andersen, 2005).

The sun emits a broad range of electromagnetic radiation, including gamma rays, X-rays, and ultraviolet, visible, and infrared light (Gueymard, 2010), which is vital to all living organisms Recently, free-electron laser (FEL) facilities have been commissioned in many places, and their radiation ranges extend from X-rays to terahertz radiation (Cohn et al., 2015; Edwards et al., 2005). Japan has LEBRA-FEL at Nihon University (Hayakawa et al., 2004; Tanaka et al., 2004) and KU-FEL at Kyoto University (Zen et al., 2013), which are outstanding facilities. These two FELs are tuneable, have high optical power and spatially coherent pulse structures. By combining the two lasers, we can provide pulsed wavelengths (1 Hz for KU-FEL, 2 Hz for LEBRA-FEL) from 1.6 to 20 μm. Harmonic generation also provides tuneable visible wavelengths up to the near-infrared region (400 nm to 2 μm).

In this study, we clarify whether the compound eyes of the crayfish (*Procambarus clarkii*) can see mid-infrared radiation, which is outside the range of human vision. A major reason why we used mid-infrared radiation is that infrared radiation (0.75–1000 μm), particularly mid-infrared radiation, can cause changes in biological molecules and biological structures (Vatansever and Hamblin, 2012), and could be a photic probe for exploring vision sensing in animals. Thus, the pulsed mid-infrared FELs generated by the two facilities were suitable for three reasons. First, an FEL can be continuously tuned within its wavelength range of operation energy easily compared with conventional lasers (Mester et al., 1985). Second, FELs in the mid-infrared region have not been used to explore vision (Edwards et al., 2005), and may promise a valuable light source for clarifying the photochemical reactions of living organisms. Third, the combination of the two FEL facilities provide visible to mid-infrared wavelengths that may be superior to other FEL facilities (Cohn et al., 2015).

Here we show that the crayfish (*Procambarus clarkii*) compound eye responds to pulsed mid-infrared FELs (1 Hz for KU-FEL; 2 Hz for LEBRA-FEL) from 3 to 17 μm. Our finding provides insight into the animal’s visual sensing of mid-infrared radiation, and that the sensing does not depend on thermal effects. Although the behavioural reason for vision in this wavelength range and the mechanism are still under investigation, understanding this type of visual sensing may lead to applications other than photophysical applications (Edwards et al., 2005).

## RESULTS AND DISCUSSION

Suction electrodes (Shishikura et al., 2015), gold wire contact electrodes (Bayer et al., 2000), and cotton wick contact electrodes (Bayer et al., 2000; Chekroud et al., 2011; Shishikura et al., 2015) were examined for recording electrical signals from crayfish eyes. The suction electrode is widely used; however, it is limited to use in physiological solution (Shishikura et al., 2015). The gold wire and cotton wick electrodes can be used in the air, and we obtained good electroretinograms (ERGs) (Brown, 1968) with high amplitudes and high signal/noise ratios from the crayfish eyes. The cotton wick electrode was used in subsequent experiments because it was used in other pioneering works by Kennedy and co-workers (Kennedy and Bruno, 1961). Hence, the cotton wick contact electrodes gave us confidence in our results, even though they have not been used to investigate vision with mid-infrared FEL radiation.

Using manipulators, a physiological saline-soaked cotton wick electrode was placed in contact with the cornea of the crayfish eye (supplementary material Fig. S1). Fig. 1A,B shows the ERGs taken for a wavelength of 5 μm at KU-FEL and LEBRA-FEL. The ERGs resemble those obtained from human eyes under scotopic (dark) conditions (Slamovits, 1993), where the a-and b-wave components are defined as follows. The a-wave is defined as the first large negative-going component, followed by the b-wave, which is generally a positive-going component. The amplitude of the a-wave is measured from the baseline to the negative trough of the a-wave, and the amplitude of the b-wave is measured from the trough of the a-wave to the following peak of the b-wave (Creel, 2017; Slamovits, 1993). Applying this definition to the crayfish compound eye, biphasic waveforms comprising a-and b-waves were detected in the ERGs generated by FEL irradiation and by LED flashes. Compared with the waveforms generated by LED flashes (61 s in, Fig. 2A), FEL irradiation generated two or three distinctive negative-going discharges with an accompanying b-wave. It is unclear why the several waves were generated by a single pulse of FEL irradiation.

**Fig. 1.**
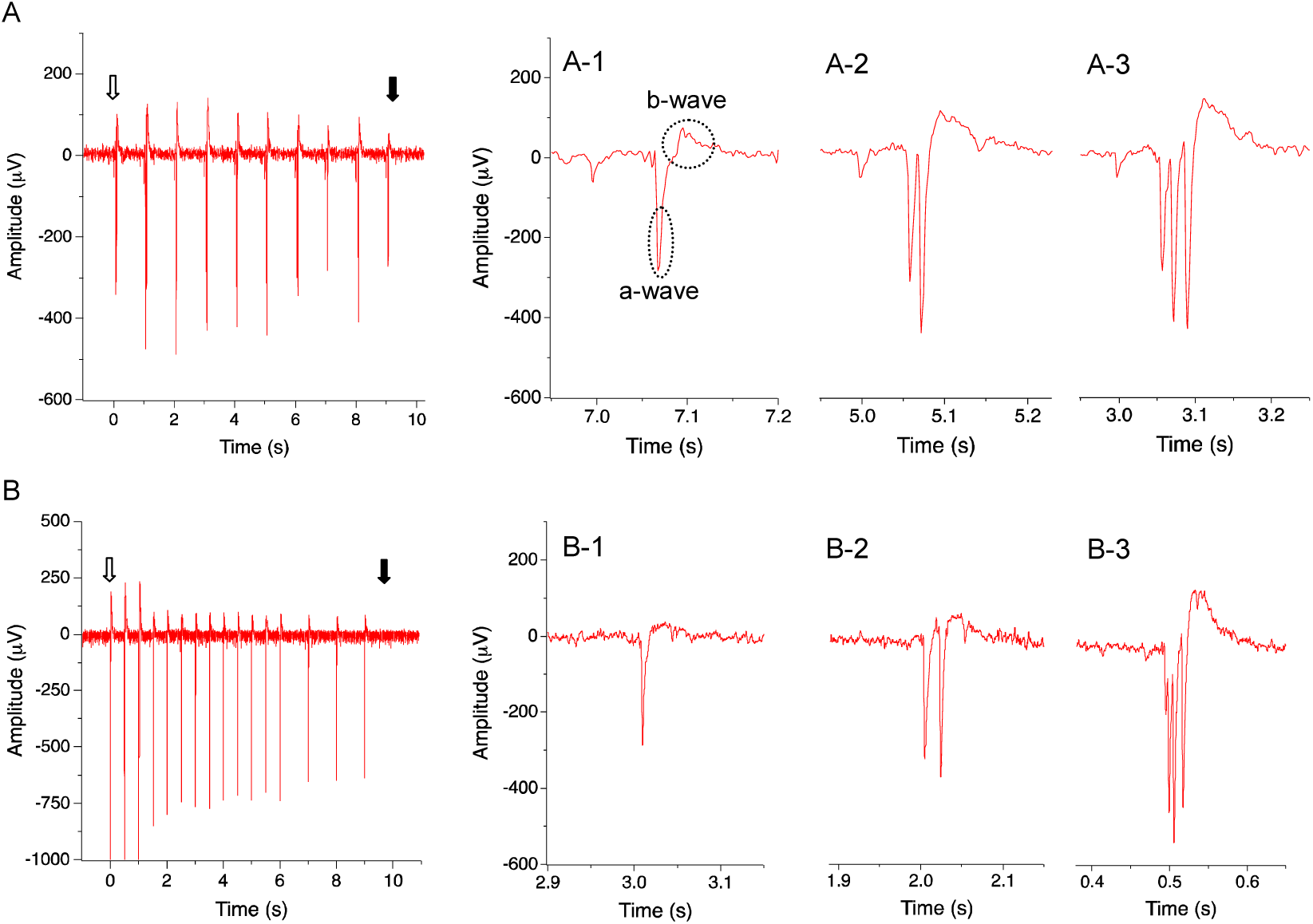
Typical discharges of the crayfish compound eye evoked by pulsed mid-infrared FEL at a wavelength of 5 μm. (A) ERGs of KU-FEL under 1 Hz irradiation (radiation intensity: 0.3 mJ/mm^2^). The three waveforms detected in A are shown in the enlarged figures: A-1, one a-wave; A-2, two a-waves; and A-3, three a-waves. The two components (a-and b-waves) are applied to A-1. (B) ERGs of LEBRA-FEL under 2 Hz irradiation (radiation intensity: 0.03 mJ/mm^2^). The waveforms resemble those of A, which may have three or four types, are shown in the enlarged figures: B-1, one a-wave, B-2, two a-waves, and B-3, four a-waves. The three enlarged figures are taken from different experiments. Two animals were used for the KU-FEL observations and four were used for the LEBRA-FEL observations. Each animal was irradiated repeatedly two to four times. There were twice as many electrical signals in B (ERGs of LEBRA-FEL) as in A over the same duration. The beginning and end of the FEL irradiation are indicated by an open arrow and a solid arrow, respectively.

**Fig. 2.**
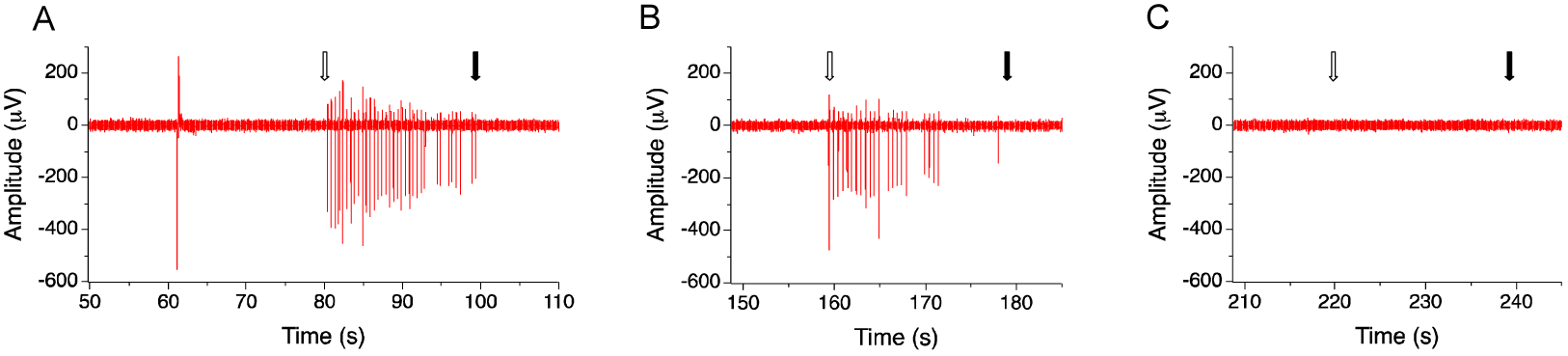
Effects of silicon and N-BK7 glass filters on mid-infrared FEL irradiation. Before LEBRA-FEL irradiation at 5 μm, a flash of LED white light was delivered as a test experiment, generating one a-wave and one accompanying b-wave (61 s in A: upper left side). (A) ERG recorded without filters. (B) Effects of the silicon filter. (C) Effects of the glass filter. The three experiments were conducted in sequence on an identical sample, the elapse times of which are shown on the time axes. The beginning and end of the FEL irradiation are indicated by an open arrow and a solid arrow, respectively.

We verified the irradiation results obtained from KU-FEL (1 Hz) facility by cross-checking with LEBRA-FEL (2 Hz). The two FEL irradiation results obtained at 5 μm (Fig. 1A,B) demonstrate that crayfish compound eyes respond to mid-infrared FEL irradiation.

Studies on infrared laser radiation in human vision have examined the foveal sensitivity to near-infrared wavelengths of 1.0 and 1.05 μm (Griffin et al., 1947) and 0.8–1.55 μm (Dmitriev et al., 1979). It has been demonstrated that human near-infrared vision is triggered by two-photon chromophore isomerization including second-harmonic generation in the eye (Palczewska et al., 2014; Zaidi and Pokorny, 1988). In this work, to avoid the effect of harmonic generation in FEL radiation, a silicon filter or an N-BK7 glass filter was placed in the FEL beam path. However, the silicon filter does not prevent the effects of harmonic radiation originating in the crayfish eyes. We predict that harmonic generation is likely to occur in our case. Therefore, we are planning to use multi-photon absorption techniques (Denk et al., 1990; Palczewska et al., 2014) to ascertain whether harmonic radiation triggered a response in our mid-infrared radiation experiments.

Fig. 2 shows the ERGs captured from the mid-infrared irradiation of LEBRA-FEL (wavelength: 5 μm) with a silicon filter or two N-BK7 glass filters. The crayfish eyes did not generate ERGs when the N-BK7 filters were inserted. In contrast, with the silicon filter, the number and the amplitude of the ERGs were reduced by more than 50% compared with the ERGs with no filter. This also suggests that crayfish eyes respond to mid-infrared wavelengths. In addition, the similarities of the ERG waveforms of the crayfish to those of humans indicate that the response of the crayfish eyes to mid-infrared radiation could be a visual reaction, not corneal injury caused by FEL radiation. Finally, we examined 18 FEL wavelengths (data not shown) of mid-infrared radiation from 3 to 20 μm, and large ERGs were obtained from 3 to 17 μm.

Nocturnal feeding animals can respond to infrared radiation emitted from warm prey animals, and they have developed various types of thermoreceptor organs known as pit organs (Campbell et al., 2002). In this work, the heat effects were negligible because both the radiation sources were continuous pulsed radiation with repetition of 1 Hz for KU-FEL and 2 Hz for LEBRA-FEL, different from continuous infrared radiation that can produce heat. The temperature was measured continuously near the surface of the eye during KU-FEL irradiation and did not increase with the irradiation over 10 s.

To reduce the FEL radiation intensity, a polarizing filter and a longwave-pass filter were placed in the FEL beam path. Fig. 3 shows the effects of the reduction of the radiation intensities for KU-FEL at a wavelength of 10 μm. The radiation intensity was changed in the sequence 0 (initial intensity: 0.27 mJ/mm^2^), 1/2 intensity, 1/4 intensity, and 1/8 intensity, and the FEL irradiation was applied for 20 s. The FEL-induced activity and post-FEL effects at different radiation intensities were recorded under dark conditions. A 1-min interval was more than sufficient for the electrical signals to return to the static background discharge, and empirically not more than a few seconds was required. At the 1/8 intensity in this typical experiment, we did not detect electrical signals except for the initial response (0 s in Fig. 3D). Therefore, the electrical discharges were a function of the light source intensity, namely the pulsed FEL radiation, and they appeared to arise from the complex reactions of the crayfish eye.

**Fig. 3.**
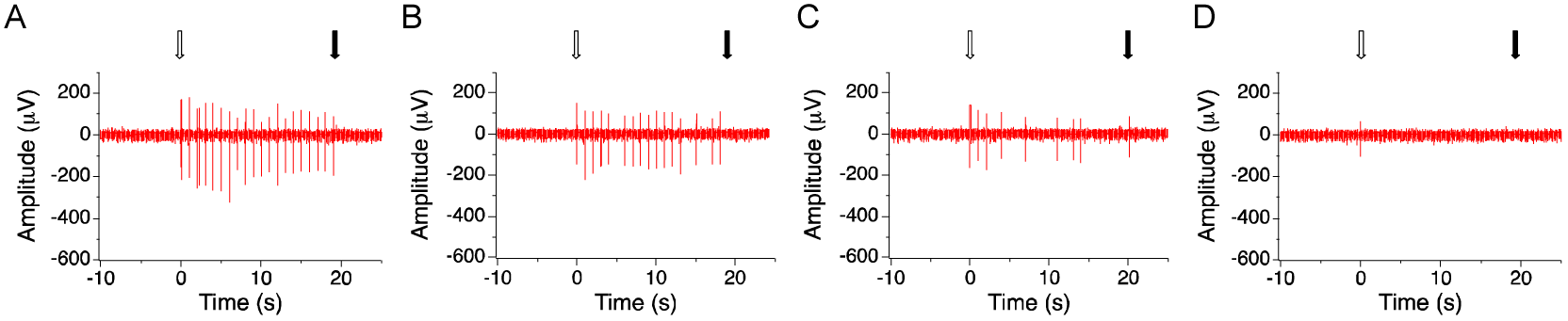
Effects of the reduction of mid-infrared FEL irradiation intensity. Before the irradiation experiments, the radiation intensity of the KU-FEL 10-μm wavelength was calibrated by using a polarizing filter with a longwave-pass filter (4.5 μm cutoff), and then the radiation intensity was fixed at 0.27 mJ/mm^2^. This value was counted as the initial radiation intensity (0-state: a polarizing angle 0). (A) Typical discharges of KU-FEL irradiation at full intensity. (B) 1/2 intensity (polarizing filter angle: 318°). (C) 1/4 intensity (polarizing filter angle: 300°). (D) 1/8 intensity (polarizing filter angle: 290°). The four experiments were conducted sequentially on the same animal. The beginning and end of the FEL irradiation are indicated by an open arrow and a solid arrow, respectively.

This study is the first to demonstrate that the compound eyes of crayfish can respond mid-infrared FEL radiation from 3 to 17 μm. We are currently clarifying the biological meaning of the mid-infrared reactions we observed in crayfish eyes.

## MATERIALS AND METHODS

### Animals

About 100 adult crayfish (*Procambarus clarkii*; body lengths, excluding the claws, of 10–11 cm) were collected from local ponds located about 70 km from LEBRA, Nihon University, and kept under a dark/light cycle of 12/12 h in separate plastic containers in fresh water about 5 cm deep that was changed on feeding. They were fed small pieces of potato and fish food twice a week. Crayfish inhabit ponds, marsh, and rivers, and, during breeding season, they are found on footpaths and in drainage ditches between rice fields, where their compound eyes are exposed to air for many hours.

### Setups of irradiation systems

The setups of the LEBRA-FEL and the KU-FEL irradiation systems have been described in previous papers (Shishikura et al., 2014; Shishikura et al., 2015). Prior to irradiation, the experimental animal was immobilized with wet paper towels and fixed with rubber bands. The compound eyes were fixed with small wet cotton wicks around the base of the eye stalks. The animal was placed in a plastic bottle to place the dorsal side and the compound eyes upward in the Faraday cage. The size of irradiation area was focused to about 5 mm in diameter by using mirrors and a lens (CaF_2_, *f*: 500 mm; Sigma-Koki, Co., Ltd., Tokyo, Japan) about 20 cm above the compound eye. Fig. S1 (supplementary material) shows a schematic diagram of the setup of the FEL irradiation system and recording apparatus.

We use the definitions in the Photonics Spectrum Reference Chart (Laurin Publishing, Pittsfield, MA) for the ranges for the visible (400–750 nm), near-infrared (750 nm–3 μm), and mid-infrared spectra (3–30 μm). Based on these criteria, we provided a full range for the visible spectrum and a major part of the mid-infrared spectrum. We focused on using mid-infrared radiation generated by the combination of the two FEL facilities, to investigate its biological effects on vision and sensing. The typical beam parameters of the two FELs have been reported elsewhere (Hayakawa et al., 2004; Tanaka et al., 2004; Zen et al., 2013). The beams had the following parameters: 1 Hz, ∼3% of the spectrum width, and a macropulse duration of 2 μs for KU-FEL; and 2 Hz, ∼0.5% of the spectrum width, and a macropulse duration of 2–20 μs for LEBRA-FEL.

The radiation intensity at the compound eye was measured by two types of power meters: pyroelectric energy detectors (PE25-BB-S and PE 10-S, Ophir Japan Ltd., Saitama City, Japan) or multi-function optical meters (818E-20-50S and 1835-C, Newport Corp., Irvine, CA). To reduce the FEL beam power, a polarizing filter (WP25H-K, Thorlabs Inc., Newton, NJ) was placed in the FEL beam path. To calibrate the radiation intensity, a collimator (2 × 2 mm) was placed in the position of the eye, and the value was fixed at 0.3–0.7 mJ/mm^2^, depending on the FEL wavelengths. Unexpectedly, we noticed that the FEL light contained small quantities of minor wavelengths generated harmonically. To remove the harmonic wavelengths, in particular the second harmonic generation (Palczewska et al., 2014; Zalidi and Pokorny, 1988), we used a silicon filter (Sigma-Koki), two longwave-pass filters (4.5 μm cutoff, No. 68-555, Edmund Optics Japan, Tokyo, Japan; 9.0 μm cutoff LP-9000 nm, Spectrogon US Inc., Mountain Lakes, NJ), and an N-BK7 glass filter (WINDOW BK7 1/4, Edmund Optics Japan; 10 mm thick).

### Recording

In a Faraday cage (supplementary material Fig. S1), a cotton wick electrode placed against the compound eye was used to record the corneal response to FEL irradiation. The wick was placed laterally on the corneal surface so that it did not interfere with the vertical FEL beam. The N-BK7 glass filter allowed wavelengths of less than 2.5 μm through and transmitted over 95% of visible light. The silicon filter transmitted approximately 50% of the infrared wavelengths from 1.2 to 6 μm, but not the visible wavelengths. The differential signal was recorded with the second electrode, a reference electrode of Ag-AgCl wire (0.25 mm in diameter; The Nilaco Co., Tokyo, Japan) that was wrapped with cotton and dampened with crayfish physiological saline (Harreveld, 1936). The reference electrode was fixed to the dorsal surface of the crayfish’s cephalothorax with a rubber band (supplementary material Fig. S1). All the procedures, except for minor changes, followed the work of Kennedy and co-workers (Kennedy and Bruno, 1961). Electrical signals (ERGs) were fed into a high-input impedance amplifier (DAM80 Differential Amplifier, World Precision Instruments, Inc., Sarasota, FL; settings: low filter, 1 Hz; high filter, 10 kHz; gain, 1000) connected to a data acquisition device (PowerLab 2/26, ADInstruments, Inc., Dunedin, New Zealand). The oscilloscope traces of ERGs were recorded and stored continuously on a computer. Recordings and irradiation experiments were performed at room temperature (15–22 °C) in a darkened room. Data analyses were performed with LabChart 7 (ADInstuments, Inc.). Prior to recording, the optical system and the light response of the compound eye were tested for reliability by using a commercially available white LED (SG-320, Gentos Ltd., Tokyo, Japan). An LED flash generated a negative electrical signal of about −500 mV (gain, 1000). If this was not observed, the measurement was repeated until acceptable electrical signals were obtained.

## Acknowledgement

This work was supported by the “Joint Usage/Research Program on Zero-Emission Energy Research”, Institute of Advanced Energy, Kyoto University ZE26C-2, ZE27C-4, and ZE28C-5).

## Author contributions

This research was conceived and designed by F.S. Experiments and measurements were executed by F.S., Z.H., K.H., and Y.K. All authors participated in the discussion, interpretation and writing process.

## Supplementary material

Supplementary material available online at http://jeb.biologists.org/lookup/suppl/doi:xxxxxxx/-/DC1

---

**Fig. S1.**
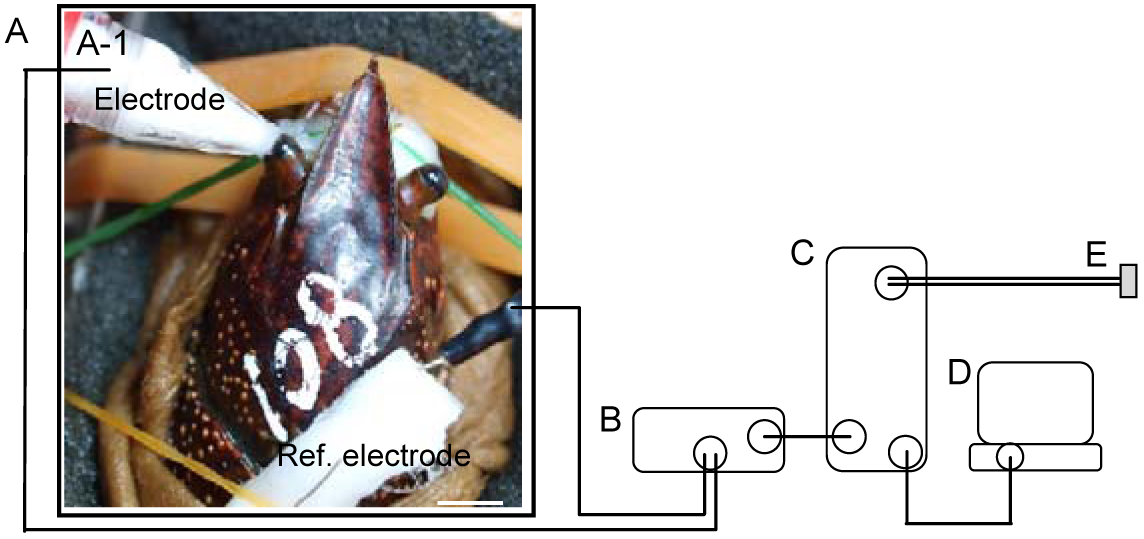
Schematic diagram showing the irradiation setup and recording apparatus for crayfish compound eyes. (A) Faraday cage. (B) DAM80 amplifier. (C) PowerLab 2/26. (D) PC with analytical software (LabChart 7 Japanese). (E) AC power supply. (A-1, inset of A) Photograph of the dorsal view of the crayfish (No. 108; adult female), showing the location of the handmade cotton wick contact electrode and the reference electrode. The bases of the eyestalks were padded with cotton wicks to immobilize the eyes. The FEL beam line came from a vertical direction to the eye, and the irradiation was focused to a 5-mm diameter spot covering the eye surface. The scale bar indicates 10 mm.

## References

Andersen, O. S. (2005). A brief history of the Journal of General Physiology. J. Gen. Physiol. 125, 3–12.

Bayer, A. U., Mittag, T., Cook, P., Brodie, S., Podos, S. M., and Maag, K.-P. (2000). Comparisons of the amplitude size and the reproducibility of three different electrodes to record the corneal flash electroretinogram in rodents. Documenta Ophthalmologica 98, 233–246.

Brown, K. T. (1968). The electroretinogram: its components and their origins. Vision Res. 8, 633–677.

Campbell, A. L., Naik, R. R., Sowards, L., and Stone, M. O. (2002). Biological infrared imaging and sensing. Micron 33, 211–225.

Chekroud, K., Arndt, C., Basset, D., Hamel, C. P., Brabet, P. and Pequignot, M. O. (2011) Simple and efficient: validation of a cotton wick electrode for animal electroretinography. Ophthalmic Res. 45, 174–179.

Cohn, K., Blau, J., Colson, W. B., Ng, J., and Price, M. (2015). Free electron lasers in 2015. Proc. FEL 2015, 625–629.

Creel, D. J. (2017). The electroreetinogram and electro-oculogram: clinical applications. In Webvision (ed. H. Kolb, R. Nelson, E. Fernandez and B. Jones) http://webvision.med.utah.edu/book/electrophysiology/the-electroretinogram-clinical-applications/

Denk, W., Strickler, J. H. Webb, W. W. (1990). Two-photon laser scanning fluorescence microscopy. Science 248: 73–76.

Dmitriev, V. G., Emelyanov, V. N., Kashintsev, M. A., Kulikov, V.V., Solovev, A. A., Stelmakh, M.F., and Cherednichenko, O. B. (1979). Nonlinear perception of infrared radiation in the 800-1355 nm range with human eye. Sov. J. Quantum Electron 9, 475–479.

Edwards, G. S., Allen, S. J., Haglund, R. F., Nemanich, R. J., Redlich, B., Simon, J. D., and Yang, W.-C. (2005). Applications of free-electron lasers in the biological and material sciences. Photochem. Photobiol. 81, 711–735.

El-Bakary, Z. A., and Sayed, A. E.-D. H. (2011). Effects of short time UV-A exposures on compound eyes and haematological parameters In Procambarus clarkii (Girard, 1852). Ecotoxicology Environmental Safety 74, 960–966.

Griffin, D. R., Hubbard, R., and Wald, G. (1947). The sensitivity of the human eye to infra-red radiation. J. Opt. Soc. Am. 37, 546–554.

Gueymard, C. (2010) Data from: SMARTS, Renewable Resource Data center, National Renewable Energy Laboratory, US Department of Energy. http://rredc.nrel.gov/solar/spectra/

Harreveld, A. van. (1936). A physiological solution for fresh-water crustaceans. Proc. Soc. Exp. Biol. Med. 34, 428–432.

Hayakawa, K., Tanaka, T., Hayakawa, Y., Yokoyama, K., Sato, I., Kanno, K., Nakao, K., Ishiwata, K., and Sakai, T. (2004). The LEBRA 125 MeV electron linac for FEL and PXR generation. Proc. LINAC 2004, 90–92.

Huxley, T. H. (1880). The Crayfish: an introduction to the study of zoology. Natural History Museum Library, London, 1880. https://archive.org/details/crayfishintroduc80huxl/

Kennedy, D., and Bruno M. S. (1961). The spectral sensitivity of crayfish and lobster vision. J. Gen. Physiol. 44, 1089–1102.

Lindstrom, M., and Meyer-Rochow, V. B. (1987). Near infra-red sensitivity of the eye of the crustacean *Mysis relicta*? Biochem. Biophys. Res. Commun. 147, 747–752.

Mester, E., Mester, A. F., and Mester, A. (1985). The biomedical effects of laser application. Lasers Surgery Med. 5, 31–39.

Nilsson, D.-E., and Kelber, A. (2007). A functional analysis of compound eye evolution. Arthropod Structure Develop. 36, 373–385.

Palczewska, G., Vinberg, F., Stremplewski, P., Bircher, M. P., Salom, D., Komar, K., Zhang, J., Cascella, M., Wojtkowski, M., Kefalov, V. J., and Palczewski, K. (2014). Human infrared vision is triggered by two-photon chromophore isomerization. PNAS published online Dec 1, 2014 E5445–E5454: www.pnas.org/cgi/doi/10.1073/pnas.1410162111.

Schmitz, H., Murtz, M., and Bleckmann, H. (2000). Responses of the infrared sensilla of *Melanophila acuminate* (Coleoptera: Buprestidae) to monochromatic infrared stimulation. J. Comp. Physiol. A 186, 543–549.

Shishikura, F., Hayakawa, K., Hayakawa, Y., Nakao, K., Inagaki, M., Nogami, K., Sakai, T, Tanaka, T., Zen, H., and Kii, T., et al. (2014). Potential photochemical applications of the free electron laser irradiation technique in living organisms. Proc. FEL 2014, 505–508.

Shishikura, F., Hayakawa, K., Hayakawa, Y., Inagaki, M., Nakao, K., Nogami, K., Sakai, T., Tanaka, T. and Komatsuzaki, Y. (2015). LEBRA free-electron laser elicits electrical spikes from the retina and optic nerve of the slug *Limax valentianus*. Proc. FEL 2015, 550–553.

Slamovits, T. L. (1993). Orbit, Eyelids, and Lacrimal System. In Basic Clinical Science Course. Section 12: San Francisco, American Academy of Ophthalmology 1993.

Tanaka, T., Hayakawa, K., Hayakawa, Y., Mori, A., Nogami, K., Sato, I., Yokoyama, K., Ishiwata, K., Kanno, K., and Nakao, K., et al. (2004). Tunability and power characteristics of the LEBRA infrared FEL. Proc. 2004 FEL Conf., 247–250.

Vatansever, F., and Hamblin, M. R. (2012). Far infrared radiation (FIR): its biological effects and medical applications. Photonics Lasers Med. 4, 255–256.

Yong, E. (2016). Seeing the light. Nat. Geographic 229, 30–57.

Zaidi, Q., and Pokorny, J. (1988). Appearance of pulsed infrared light: second harmonic generation in the eye. Applied Optics 27, 1064–1068.

Zen, H., Inukai, M., Okumura, K., Mishima, K., Torgasin, K., Negm, H., Omer, M., Yoshida, K., Kinjo, R., and Kii, T., et al. (2013). Present status of Kyoto University free electron laser. Proc. FEL 2013, 711–714.

